# Intrinsic Hydrogen Deuterium Exchange Rates in H_2_O/D_2_O Mixtures

**DOI:** 10.1101/2025.09.11.674334

**Authors:** Antonio Grimaldi, Michele Stofella, Emanuele Paci

## Abstract

Hydrogen–deuterium exchange (HDX) measurements are widely used to probe protein structural dynamics. Quantitative interpretation of HDX data relies on the concept of an intrinsic exchange rate, which is well characterized in isotopically pure H_2_O or D_2_O but does not explicitly account for the back exchange that necessarily occurs in H_2_O/D_2_O mixtures: in this case, both the approach-to-equilibrium rate and the equilibrium deuterium enrichment of amides depend nontrivially on solvent composition and acidity. A practical method is presented to predict intrinsic forward and reverse amide exchange rates in H_2_O/D_2_O mixtures. The approach combines known second-order reference rates measured in pure solvents with established empirical descriptions of H_2_O/D_2_O mixtures. The resulting framework yields explicit expressions for forward and back exchange rates as functions of solvent composition and acidity, and correctly recovers the known limits in pure H_2_O and pure D_2_O. The model predicts composition-dependent kinetic isotope effects and an equilibrium amide fractionation factor of *ϕ* = 1.20 for unstructured peptides in base-catalyzed conditions, in close agreement with the experimental value 1.22 reported for poly-D,L-alanine. By providing a physically motivated description of exchange in mixed solvents, this method offers a practical starting point for quantitatively correcting back exchange in HDX–MS and HDX–NMR experiments.

## Introduction

Biophysical techniques measuring amide hydrogen-deuterium exchange (HDX) are broadly used to fingerprint structural dynamics of proteins.^1–4^ The Linderstrøm-Lang model describes HDX of protein backbone amides according to the reaction^5^

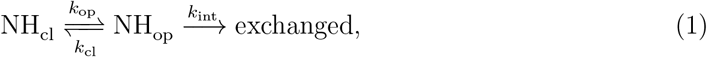

where the amide can assume either a closed (NH_cl_) or open (NH_op_) conformation, the former being exchange-incompetent. An amide in the open state exchanges with a pseudo-first-order rate *k*_int_, called the intrinsic (chemical) exchange rate, which can be estimated as a function of sequence, temperature and pH (or pD) in pure H_2_O (or D_2_O).^6–8^ The protection factor *P* = *k*_cl_*/k*_op_ is the equilibrium constant for site closing ^3^ that one aims to determine to characterize structural dynamics.^9–14^ In EX2 conditions (*k*_cl_ ≫ *k*_op_, *k*_int_), the protection factor is related to the observed exchange rate *k*_obs_ as *P* = *k*_int_*/k*_obs_.

HDX is typically detected using nuclear magnetic resonance (NMR) spectroscopy or mass spectrometry (MS). NMR experiments and the labeling step in HDX-MS are commonly conducted in buffers containing 80-95% D_2_O.^15^ While mixed-solvent equilibrium and kinetic isotope effects are well established, ^16,17^ they are often treated approximately or neglected in practical analyses, despite a body of work that used HDX in H_2_O/D_2_O mixtures to quantify amide fractionation and probe hydrogen-bonded structure. ^18–21^ Mixed H_2_O/D_2_O solvents are also deliberately used in NMR protocols that quantify amide–solvent exchange, including CLEANEX-PM^22^ and SOLEXSY.^23^

In H_2_O/D_2_O mixtures, exchange is intrinsically bidirectional and cannot be described by a single rate constant. n this work, we make this explicit by decomposing intrinsic exchange into forward and reverse components, enabling a quantitative treatment of kinetic and equilibrium isotope effects in mixed solvents that is not accessible within standard intrinsic-rate models. This work presents a practical method to predict forward and reverse amide exchange rates in H_2_O/D_2_O mixtures as functions of the degree of deuteration and acidity. The approach combines published reference parameters measured in pure solvents with an operational acidity scale for mixtures, and a probabilistic treatment for reprotonation inserting H or D. The resulting expressions recover pure-solvent limits and provide explicit mixture-dependent rates.

## Methods

### Proton transfer theory

The chemistry of HDX reactions is described by proton transfer theory:^3,24^

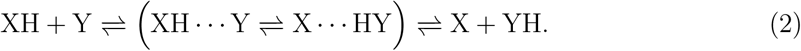

A proton donor XH collides with a proton acceptor Y, forming a short-lived encounter complex (bracketed in Eq. 2) in which proton redistribution is faster than dissociation. Here, XH and Y are generic acid/base labels and may be neutral or charged depending on the specific mechanism, the corresponding conjugate-acid pairs being XH/X and YH/Y. Complex formation occurs with a second-order (diffusion-limited) rate constant *k*_d_ estimated as 10^10^ M^−1^ s^−1^.^25^ The complex is regarded as highly dynamical but always weakly populated. Focusing on the forward process XH + Y →X + HY, the proton transfer rate constant *k*_t_ can be expressed as^3^

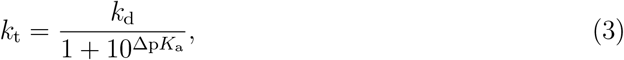

where Δp*K*_a_ = p*K*_a_(XH) − p*K*_a_(YH) is the difference between the acidity of the donor XH and that of the conjugate acid YH of the acceptor. The acidity constant of a generic acid HA is defined as

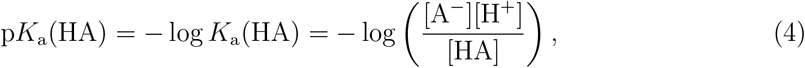

with *K*_a_(HA) being the acid dissociation constant.

### Intrinsic exchange rates

In 100% D_2_O, amide HDX involves removal of a proton (H^+^) from the amide group and the transfer of a deuteron (D^+^) from bulk solvent to the amide. The reaction can be acid-, base-, or water-catalyzed, and the pseudo-first-order HDX intrinsic exchange rate of Eq. 1 can be written as

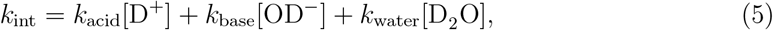

where *k*_acid_, *k*_base_ and *k*_water_ are the corresponding second-order rate constants. Temperature and pH dependence of HDX second-order rate constants were measured by Englander’s group for poly-D,L-alanine (PDLA) and 3-alanine (3-Ala). ^6–8^

Below, we dissect the base-catalyzed mechanism of HDX, which dominates by orders of magnitude at near-neutral conditions.^3^ Extension to other mechanisms (acid- and water-catalyzed) is reported in the Appendix. For base-catalyzed HDX in pure D_2_O, the kinetic bottleneck is the first step, i.e. abstraction of the amide proton by OD^−^. The amide is rapidly deuterated by a solvent D_2_O molecule, and a new OD^−^ ion is produced. In this case, the intrinsic rate can be estimated as

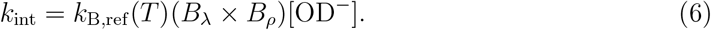

*B*_*λ*_ and *B*_*ρ*_ are tabulated factors that depend on amino acid type and position relative to the amide group (*λ* for left, *ρ* for right, see Table A1).^6^ These account for steric and inductive effects from neighboring residues, and were demonstrated to be simply additive. ^26^ *k*_B,ref_ (*T*) is the reference rate (i.e. the exchange rate for PDLA or 3-Ala) at temperature *T*, which exhibits an Arrhenius-type dependence:

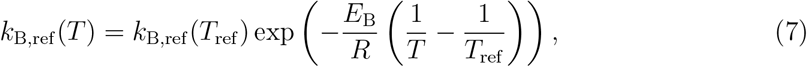

where *E*_B_ = 17 kcal mol^−1^, *R* is the gas constant, and *k*_B,ref_ (*T*_ref_) is the second-order rate constant measured. Values for *k*_B,ref_ (20°C) for H and D in H_2_O and H in D_2_O are given in Table 1.^6–8^

**Table 1:**
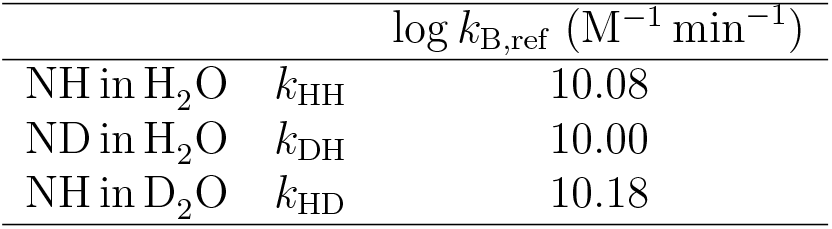
Reference rates (PDLA) for the base-catalyzed exchange of amides in disordered peptides, at 20°C.^8^ Reference rates are specified by two pedices, the first indicating the amide-bound isotope being removed, the second indicating the isotope in the catalytic base abstracting the proton, *cfr Forward and back exchange rates* Section. Rates for 3-Ala are obtained multiplying the reference rates by a factor 1.35.^8^

### H_2_O/D_2_O mixtures

This section introduces an operational acidity scale for mixed H_2_O/D_2_O solvents and then analyzes the influence of solvent composition on hydroxide availability and reprotonation probabilities that dictate intrinsic exchange rates. Let L denote either protium (H) or deuterium (D) isotope. An isotopic exchange reaction in a solvent with D_2_O mole fraction *x* (H_2_O mole fraction 1 − *x*) involving a solute XL is

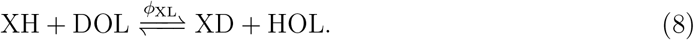

Here, XH and XD are the solute protiated and deuterated species, while HOL and DOL are solvent molecules containing at least one H or D atom, respectively. The equilibrium constant *ϕ*_XL_ of the reaction (8) is called the fractionation factor of XL

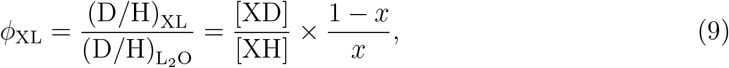

which quantifies the enrichment in D of solute XL with respect to the solvent. ^16^

The concentrations [H^+^] and [OH^−^] in pure H_2_O follow from the pH and the ionic product, 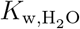, through

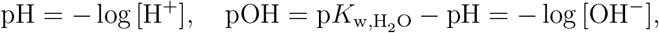

Where 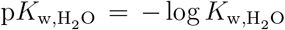. Analogous relations hold in pure D_2_O defining [D^+^] and [OD^−^] in terms of pD, pOD and 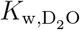.

In H_2_O/D_2_O mixtures, multiple isotopologue ionization equilibria coexist (involving H_2_O, HDO, D_2_O and the corresponding ions). Rather than assigning separate equilibrium constants to each elementary process, it is convenient to work with an operational acidity scale based on the total concentration of [L^+^] = [H^+^] + [D^+^], introducing the pL

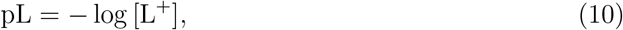

as used in studies of mixed H_2_O/D_2_O solvents by glass electrodes. ^27^ Similarly one can define the pOL via

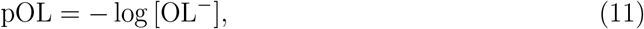

where [OL^−^] = [OH^−^] + [OD^−^]. At 25°C, the ionic product of D_2_O 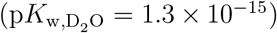 is lower than that of H_2_O 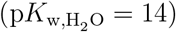.^28,29^ In mixed solvents, the corresponding effective ionic product pK_w_(*x*) = pL(*x*) +pOL(*x*) varies nonlinearly with the deuterium mole fraction *x*, and can be described empirically as ^30^

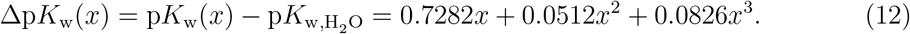

To relate the mixture pL(*x*) to the glass electrode pH-meter reading, pH^∗^, the following relation can be used:^31^

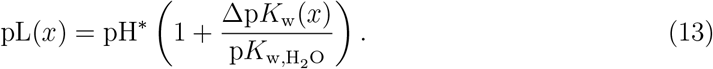

For D_2_O at 25°C and pH^∗^ = 7, Eq. 13 yields pD = pL(1) ≃ 7.43, i.e., the traditional “+0.4” correction widely used by HDX practitioners.

The temperature dependence of 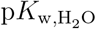 has been empirically determined as^32^

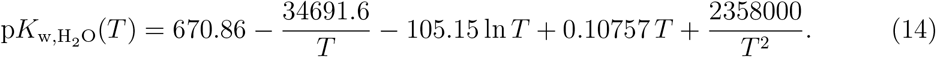

At given composition *x*, the p*K*(*x*) can be computed via Eq. 12. Once the mixture pH^∗^ is measured, the pL(*x*) can be computed using Eq. 13 and the pOL(*x*) is given by subtraction of the two values as stated above. Reported values for the fractionation factor *ϕ*_OL_ (0.47^31^ or 0.43^33^) can be used to compute [OH^−^](*x*) and [OD^−^](*x*), *cfr* Eq. A4. Examples are shown in the upper and middle panels of Figure 1.

**Figure 1:**
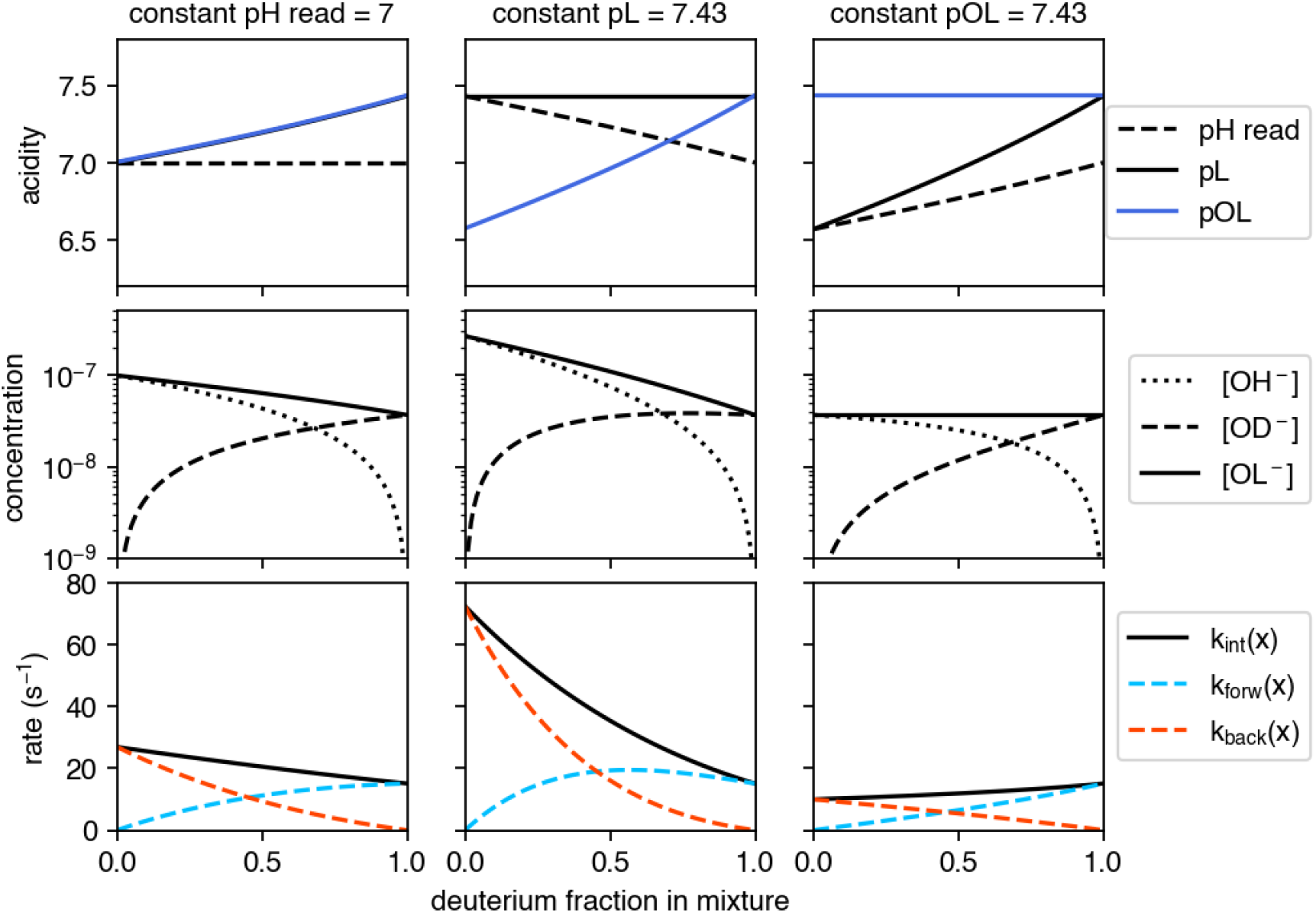
Acidity, concentration of ions and intrinsic rates in mixtures, expressed as a function of the deuterium fraction in the mixture, in three different scenarios: constant pH read (left), constant pL (center), constant pOL (right). Upper panels: mixture pH read, pL and pOL. Middle panels: concentrations of the ions relevant for base-catalyzed HDX: [OH^−^], [OD^−^], and [OL^−^] = [OH^−^] + [OD^−^]. Lower panels: intrinsic forward *k*_forw_, back *k*_back_ and mixture intrinsic *k*_int_(*x*) = *k*_forw_(*x*) + *k*_back_(*x*) exchange rates of PDLA. Magnitude and direction of kinetic isotope effects depend on the quantity fixed while exploring different H_2_O/D_2_O compositions. Data for 25°C.

### Forward and back exchange rates

The reaction considered for HDX of an open amide in a H_2_O/D_2_O mixture is

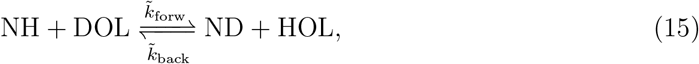

where 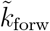 and 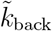 are the base-catalyzed reactions rate constants. Reference rates in pure solvents (Table 1) are denoted *k*_HH_, *k*_DH_, *k*_HD_, *k*_DD_ respectively. Here, the first index indicates the isotope initially bound to the amide (NH or ND) and the second indicates the isotope of the catalytic base (OH^−^ or OD^−^) performing proton abstraction. These correspond to the base-catalyzed rate removal of the amide-bound proton, followed by rapid reprotonation from the bulk solvent. The unmeasured rate *k*_DD_ was estimated as

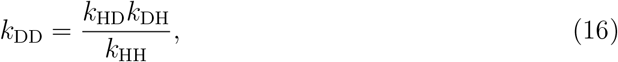

that is, assuming that reaction rates are governed entirely by the zero-point energies of the participating species.^34^

The rates of reaction (15) are obtained as a sum of terms, *cfr* Eq. 5, each having the same structure as in Eq. 6:

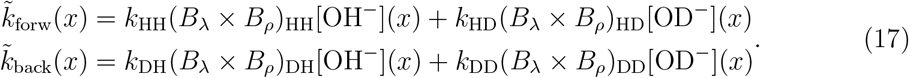

The coefficients of the products (*B*_*λ*_ *× B*_*ρ*_) are tabulated, *cfr* Appendix.

The scheme (15) can be simplified to a pseudo-first order reaction

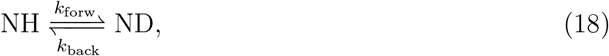

where the pseudo-first order forward and back exchange rates are weighted by the probability of encountering a reactive solvent molecule:

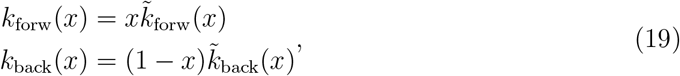

that is, upon removal of the proton, in the forward reaction reprotonation occurs by DOL with probability *x* = [DOL]*/*[L_2_O], and by HOL with probability 1 − *x* = [HOL]*/*[L_2_O] in the reverse.

The solution to the kinetics in Eq. 18, with the constraint d ([NH] + [ND]) */*d*t* = 0 and calling *D*(*t*) the normalized concentration of deuterated amides over time, is

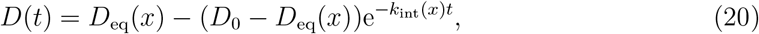

where *D*_0_ is the initial condition,

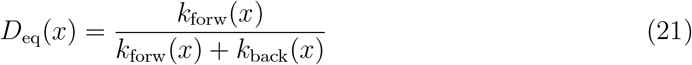

is the fraction of deuterated amides at equilibrium, and the intrinsic rate in the mixture is defined as

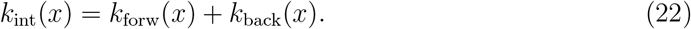

## Results and discussion

The rates *k*_forw_ and *k*_back_ are generally different and depend on the fraction *x* of D_2_O in the solvent. The stationary state (Eq. 21), as well as the approach-to-equilibrium rate (Eq. 22), are functions of *x* through *k*_forw_(*x*) and *k*_back_(*x*), *cfr* Eq. 19. In other words, kinetic and equilibrium isotope effects are present.

### Kinetic isotope effects

Kinetic isotope effects cause the reaction rate *k*_int_(*x*) to vary as a function of the solvent composition. These are due not only to the different second order rates of Table 1, but also to the solvent composition *x*, which influences the mixture ion product p*K*_w_(*x*) and acidity pL(*x*), *cfr* Eqs. 12 and 13.

By construction, the measured reference rates are recovered in pure H_2_O and D_2_O: by Eqs. 17 and 19, for *x* = 0 (pure H_2_O),

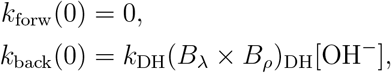

while for *x* = 1 (pure D_2_O),

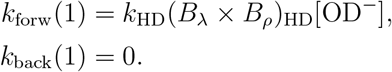

The question “how does *k*_int_(*x*) vary with *x*?” has no unique answer: it depends on the acidity measure held constant while varying *x*. At fixed temperature, the ionic product of the mixture p*K*_w_(*x*) is assumed to depend on *x* only, *cfr* Eq. 12. The observed exchange rate depends on the acidity of the solution through the concentration of ions OH^−^ and OD^−^, *cfr* Eq. 17. However, different definitions of acidity are possible in a mixture, e.g. based on pH^∗^, pL or pOL. The behavior of *k*_int_(*x*) in these scenarios is illustrated in the lower panels of Figure 1 and commented below. These results extend the observations made for pure solvents^7^ to mixtures.

In the first column of Figure 1, mixtures of different compositions that yield same pH^∗^ are considered. Starting from neutral pure H_2_O at 25°C (pH = 7), for *x* > 0 one finds 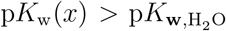 (Eq. 12). For varying composition, the solution remains neutral, thus both pL and pOL increase. A higher pOL implies fewer available OL^−^ ions for base-catalyzed HDX, hence *k*_int_(*x*) decreases for increasing *x*. As a result, the intrinsic rate of H → D in pure D_2_O is about 2-fold slower than D → H in pure H_2_O.

In the second example, the effective acidity of the mixture, pL, was fixed to 7.43 (because pL(1) = 7.43 at pH^∗^ = 7 and *T* = 25°C). Here, decreasing *x* causes p*K*_w_(*x*) to decrease. Because pL is fixed, pOL decreases too, and a more substantial kinetic isotope effect is observed. D → H in H_2_O results about 5-fold faster than H → D in D_2_O on a pL scale. The third scenario evaluates the effect of *x* at fixed pOL. In this case, *k*_int_(*x*) increases with *x*. This is because the concentration of catalysts is fixed hence the HDX rates depends on the rates of Table 1. Since *k*_HH_ > *k*_DH_ and *k*_HD_ > *k*_DD_, that is, it is easier to extract H than D from the amide group, the HDX rate is higher for increasing D_2_O content. In this case, the extent of the isotope effect (moving from pure D_2_O to H_2_O) is smaller than the other cases (1.5 times faster in D_2_O).

As an example, Figure 2 displays the approach-to-equilibrium rates *k*_int_(*x*) (Eq. 22) for two short sequences at fixed pH^∗^, which is the experimentally controlled parameter, and varying D_2_O content in the mixture. Kinetic isotope effects lead to a decrease of the reaction rate constant when the D_2_O content is increased, as discussed above.

**Figure 2:**
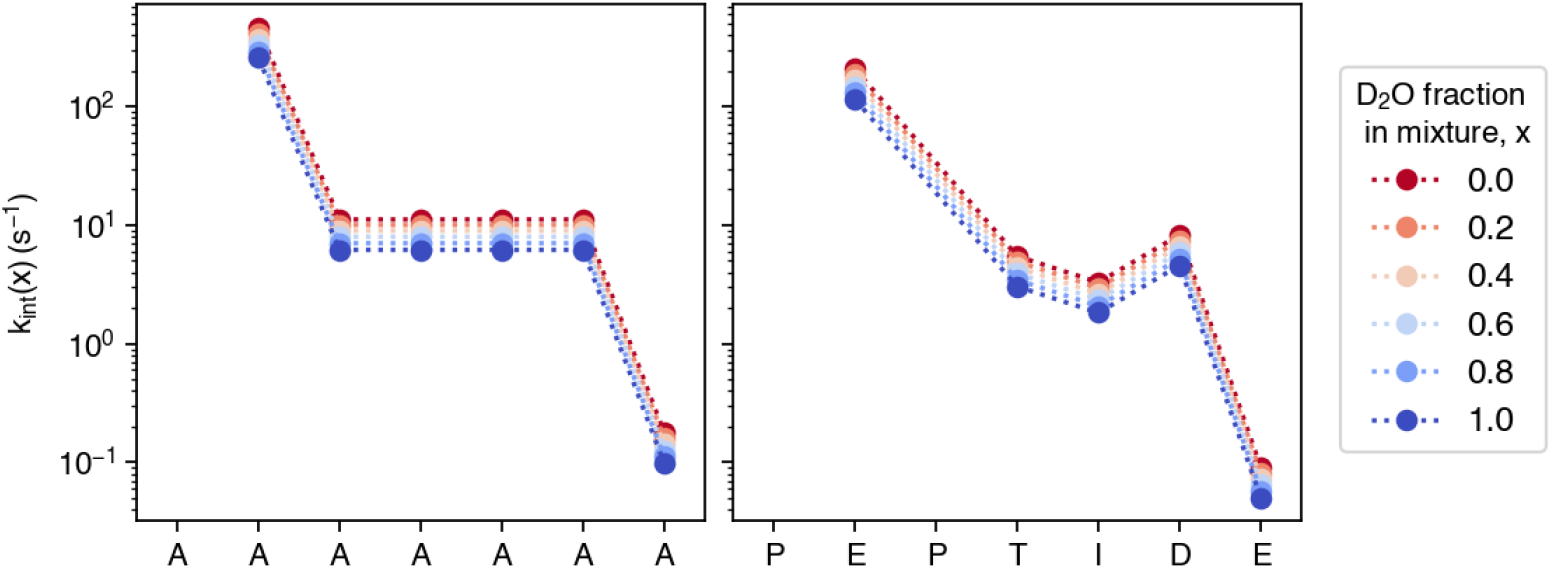
Intrinsic exchange rates *k*_int_(*x*) for short polypeptide sequences ‘`AAAAAAA`’ (left) and ‘`PEPTIDE`’ (right), calculated for varying D_2_O fraction in the mixture *x, T* = 25°C, pH^∗^ = 7, and using PDLA reference rates.

### Equilibrium isotope effects

An equilibrium isotope effect is described by the fractionation factor (Eq. 9), which for reaction (15) is simply 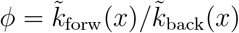. Within the assumptions of the base-catalyzed model developed above, the fractionation factor reduces to

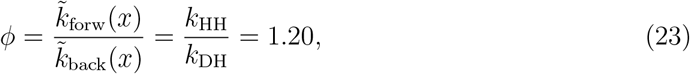

with *k*_HH_ and *k*_DH_ from Table 1.

This value is unaffected by neighboring side chains, because the local sequence multiplicative factors (*B*_*λ*_ *× B*_*ρ*_) cancel in the ratio, *cfr* Appendix. A corollary of Eq. 23 is that, for random coil peptides in conditions where base catalysis dominates, the equilibrium deuterium enrichment of amides will exceed the bulk D_2_O content, i.e. *D*_eq_(*x*) > *x*. This is shown in Figure 3. A physical basis of this difference is given by the difference in the p*K*_a_ between amide deuterated (ND) and protiated (NH) forms: p*K*_a_(ND) *>* p*K*_a_(NH), thus D tends to be bound more tightly to the amide, and the amide is enriched with D at equilibrium.^7^ The value *ϕ* = 1.20, as well as the predicted deuterium enrichment, is insensitive to temperature and pH (provided that the base-catalyzed mechanism is the dominant one).

**Figure 3:**
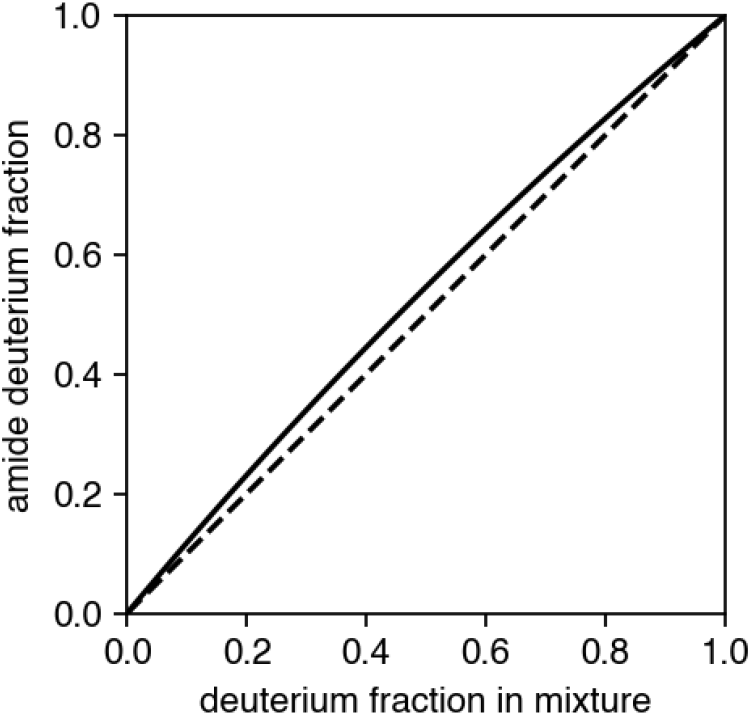
Equilibrium isotope effects. At equilibrium, the amide is enriched in D with respect to solvent (*D*_eq_(*x*) > *x*, as *ϕ* = 1.20 *>* 1).

The theoretical value *ϕ* = 1.20 (Eq. 23) agrees closely with the measured one for PDLA (*ϕ*_PDLA_ = 1.22).^19^ The larger fractionation factor reported for poly-D,L-lysine (PDLK) within the same study ^19^ (*ϕ*_PDLK_ = 1.46) may be attributed to a residual helical propensity of PDLA.^35^ It is thus possible that the equilibrium fractionation expected for unstructured peptides is underestimated, since structured amides generally display lower fractionation (usually, *ϕ* ≲ 1) than unstructured ones.^18,19^ Residual uncertainties remain also because acid- and water-catalyzed exchange, solvent–side-chain interactions, and buffer composition can all shift the apparent fractionation. Notably, the equilibrium fractionation factor emerges directly from the kinetic framework without introducing additional adjustable parameters or sequence-specific assumptions. This indicates that the observed fractionation is an inherent consequence of intrinsic exchange chemistry in mixed solvents rather than an independent thermodynamic input.

## Conclusions

This work provides a practical method to estimate intrinsic exchange rates in H_2_O/D_2_O mixtures as a function of solvent composition and acidity, recovering known limits for pure solvents. ^6–8^ The model yields closed expressions for forward and back exchange rates, that for base-catalyzed exchange incorporate mixture-dependent catalyst availability and probabilistic insertion of H or D. For completeness, the corresponding acid- and water-catalyzed contributions are described in the Appendix.

Kinetic and equilibrium isotope effects emerge as natural consequences of the chemistry of the mixture and are not determined by composition alone: both the magnitude and the direction of the kinetic isotope effect depend on how acidity is specified or experimentally controlled (e.g. fixed pH^∗^ versus fixed operational acidity). Under base-catalyzed conditions, the framework predicts equilibrium amide enrichment in D, in good agreement with measurements performed on PDLA (*ϕ* ≃ 1.20).^19^ Mismatch with PDLK (*ϕ* ≃ 1.46)^19^ suggests that residual structure or other unmodeled factors can affect fractionation and should be assessed by systematic benchmarking against random-coil standards.

Beyond providing explicit intrinsic exchange rates in mixtures, this framework emphasizes that HDX is bidirectional and that should be described by both forward and back exchange rates to capture kinetic and equilibrium isotope effects. The same framework can be applied to HDX-NMR data, where mixture-dependent kinetics and final enrichment must be accounted for to obtain unbiased estimates of local stability (in terms of protection factors). ^36^ This is also relevant to HDX-MS workflows, where back exchange can occur both during labeling (due to incomplete buffer deuteration) and in subsequent handling steps (e.g. quenching, usually performed at intermediate D_2_O content, low temperature and pH). ^15^ A consistent treatment of exchange in mixture is the first step towards physically-grounded back exchange corrections. In this case, the chemistry of the mixture defines baseline effects, that must be integrated with the environment (e.g. salts, co-solvents, denaturants, and whatever additive that can alter exchange kinetics).

## Code availability

A Python^37^ implementation of the method presented, allowing to compute *k*_forw_ and *k*_back_ for arbitrary sequences as a function of solvent temperature, glass electrode pH read, and composition, is available at https://github.com/pacilab/hdx-rates-mixtures.

## Acknowledgement

AG was supported by the Italian Ministry of University and Research (MUR) under the National Recovery and Resilience Plan (PNRR), through a PhD scholarship at the University of Bologna (DM 118/2023, CUP J33C23002540002).

## Appendix

### Multiplicative factors for intrinsic exchange rates

**Table A1:** Multiplicative factors for intrinsic exchange rates. ^6–8^.

| | $\log A_\lambda$ | $\log A_\rho$ | $\log B_\lambda$ | $\log B_\rho$ |
| --- | --- | --- | --- | --- |
| A | 0.00 | 0.00 | 0.00 | 0.00 |
| C | -0.54 | -0.46 | 0.62 | 0.55 |
| C2 | -0.74 | -0.58 | 0.55 | 0.46 |
| D0 | 0.90 | 0.58 | 0.10 | -0.18 |
| D+ | -0.90 | -0.12 | 0.69 | 0.60 |
| E0 | -0.90 | 0.31 | -0.11 | -0.15 |
| E+ | -0.60 | -0.27 | 0.24 | 0.39 |
| F | -0.52 | -0.43 | -0.24 | 0.06 |
| G | -0.22 | 0.22 | -0.03 | 0.17 |
| HO | — | — | -0.10 | 0.14 |
| H+ | -0.80 | -0.51 | 0.80 | 0.83 |
| I | -0.91 | -0.59 | -0.73 | -0.23 |
| K | -0.56 | -0.29 | -0.04 | 0.12 |
| L | -0.57 | -0.13 | -0.58 | -0.21 |
| M | -0.64 | -0.28 | -0.01 | 0.11 |
| N | -0.58 | -0.13 | 0.49 | 0.32 |
| P | — | -0.19 | — | -0.24 |
| Pc | — | -0.85 | — | 0.60 |
| Q | -0.47 | -0.27 | 0.06 | 0.20 |
| R | -0.59 | -0.32 | 0.08 | 0.22 |
| S | -0.44 | -0.39 | 0.37 | 0.30 |
| T | -0.79 | -0.47 | -0.07 | 0.20 |
| V | -0.74 | -0.30 | -0.70 | -0.14 |
| W | -0.40 | -0.44 | -0.41 | -0.11 |
| Y | -0.41 | -0.37 | -0.27 | 0.05 |
| NT | — | -1.32 | — | 1.62 |
| CT | 0.96 | — | -1.80 | — |

Coefficients *A*_*λ*_, *A*_*ρ*_, *B*_*λ*_, *B*_*ρ*_ are given in Table A1 in both H_2_O and D_2_O for all residues except Asp (`D`), Glu (`E`) and His (`H`).^8^ In these cases, the coefficients *A*_*ζ*,L_, *B*_*ζ*,L_(`X`) of residue `X`, where *ζ* is *λ* or *ρ*, and L is H (for D → H exchange) or D (H → D), are given by

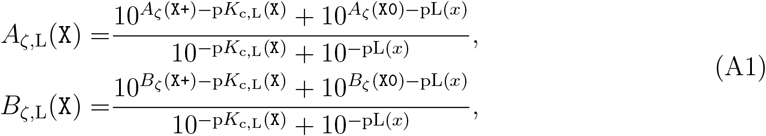

where

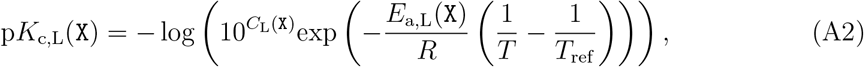

and parameters *C*_L_(`X`), *E*_a,L_(`X`), *T*_ref_ given in Table A2.

**Table A2:** Parameters for the computation of *B*_*λ*_ and *B*_*ρ*_ coefficients for Asp, Glu, His. ^8^.

| residue | $E_{a,H}$ (kcal/mol) | $C_H$ | $E_{a,D}$ (kcal/mol) | $C_D$ | $T_{\text{ref}}$(K) |
| --- | --- | --- | --- | --- | --- |
| D | 0.96 | -3.87 | 1 | -4.48 | 278 |
| E | 1.083 | -4.33 | 1.083 | -4.93 | 278 |
| H | 7.5 | -7 | 7.5 | -7.42 | 278 |

### Calculation of *ϕ*

From definitions of Eqs. A1 and A2, one has that (*B*_*λ*_ *× B*_*ρ*_)_HH_ = (*B*_*λ*_ *× B*_*ρ*_)_DH_ and (*B*_*λ*_ *× B*_*ρ*_)_DD_ = (*B*_*λ*_ *× B*_*ρ*_)_HD_. Calling *ξ* = (*B*_*λ*_ *× B*_*ρ*_)_HD_*/*(*B*_*λ*_ *× B*_*ρ*_)_HH_ (*ξ* = 1 if Asp, Glu or His are not neighboring to the amide group), the rates 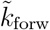 and 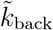 of Eq. 17 are written

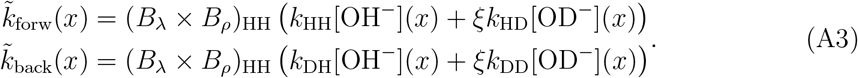

[OH^−^](*x*) and [OD^−^](*x*) can be written in terms of [OL^−^](*x*) using the definition of fractional abundance of OD^−^ in OL^−^ :^16^

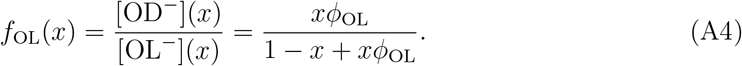

Hence Eq. A3 becomes

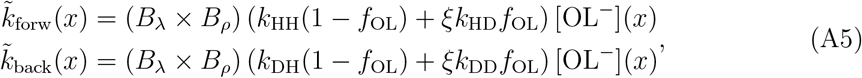

where the subscript “HH” of (*B*_*λ*_ *× B*_*ρ*_) has been dropped to ease the notation. Recalling Eq. 16, and defining *ψ* = *k*_HD_*/k*_HH_ = *k*_DD_*/k*_DH_,

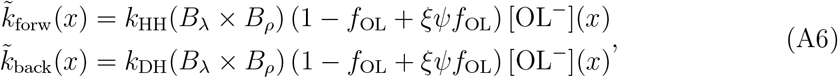

from which one has

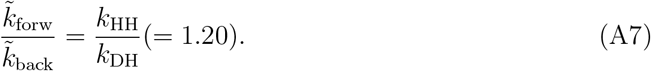

### Extension to other mechanisms

In H_2_O/D_2_O mixtures (D_2_O mole fraction *x*), the intrinsic exchange rates *k*_forw_(*x*), *k*_back_(*x*) are given as a sum of contributions (of acid-, base-, and water-catalyzed exchange):

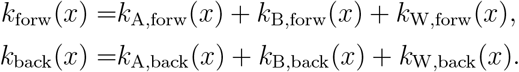

In the main text, the derivation focused on the base-catalyzed contribution, which dominates near-neutral conditions and therefore governs most HDX measurements. For completeness, this appendix outlines how the other two pathways can be incorporated into the same framework as presented in details in the main text for the case of the base-catalyzed pathway.

**Table A3:** Reference rates for acid- and water-catalyzed exchange of at 20°C.^8^.

| | | $\log k_{\text{A,ref}} \text{ (M}^{-1} \text{ min}^{-1}\text{)}$ | | $\log k_{\text{W,ref}} \text{ (M}^{-1} \text{ min}^{-1}\text{)}$ |
| --- | --- | --- | --- | --- |
| NH in D<sub>2</sub>O | $k_{\text{A}}^{\text{H} \rightarrow \text{D}}$ | 1.62 | $k_{\text{W}}^{\text{H} \rightarrow \text{D}}$ | -1.50 |
| ND in H<sub>2</sub>O | $k_{\text{A}}^{\text{D} \rightarrow \text{H}}$ | 1.40 | $k_{\text{W}}^{\text{D} \rightarrow \text{H}}$ | -1.60 |

#### Water-catalyzed exchange

In the water-catalyzed pathway, a solvent water molecule acts as a base, abstracting the amide H or D. This contribution is essentially pH-independent and usually negligible compared to the other terms, except in the region of the minimum. Sequence effects are addressed by the coefficients of Table A1, as *k*_W_ = *k*_W,ref_ (*B*_*λ*_ *× B*_*ρ*_), with reference rates given in Table A3. Literature tabulates the rates of H → D exchange in D_2_O 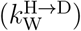 and the rate of D → H in H_2_O 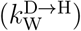. In a mixed solvent with deuterium mole fraction *x*, it is assumed that the rate of abstraction depends only on the isotope being removed, while the isotope incorporated upon reprotonation is determined solely by the solvent composition. In this case, the forward and back water-catalyzed intrinsic exchange rates are

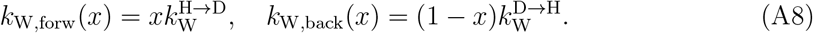

#### Acid-catalyzed exchange

In the acid-catalyzed pathway, exchange proceeds through protonation of the amide by solvated hydronium species before loss of the bound isotope. In this case, the isotopic identity of the proton-donating species, i.e. D^+^ or H^+^, determines whether the event contributes to H → D or D → H exchange. Acid-catalyzed second-order rates 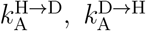 can be calculated from reference values (Table A3) and accounting for the sequence dependence with the corresponding multiplicative factors *A*_*λ*_, *A*_*ρ*_ from Table A1, e.g. 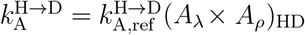. Since the key isotopic selectivity lies in the isotope donated to the amide, it is assumed that the intrinsic rate remains the same as in pure solvent, but the relevant concentrations [H^+^](*x*) and [D^+^](*x*) should be used:

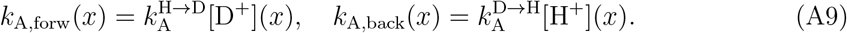

The relative abundance of [D^+^] is given, analogously to Eq. A4, as

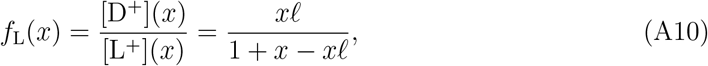

where *ℓ* = 0.69^16^ is the fractionation parameter for L_3_O^+^ and [L^+^](*x*) = [H^+^](*x*) + [D^+^](*x*), *cfr* main text.

## References

(1) Masson, G. R.; Burke, J. E.; Ahn, N. G.; Anand, G. S.; Borchers, C.; Brier, S.; Bou-Assaf, G. M.; Engen, J. R.; Englander, S. W.; Faber, J.; others Recommendations for performing, interpreting and reporting hydrogen deuterium exchange mass spectrometry (HDX-MS) experiments. Nature methods 2019, 16, 595–602.

(2) Narang, D.; Lento, C.; J. Wilson, D. HDX-MS: an analytical tool to capture protein motion in action. Biomedicines 2020, 8, 224.

(3) Hamuro, Y. Tutorial: Chemistry of Hydrogen/Deuterium Exchange Mass Spectrometry. Journal of the American Society for Mass Spectrometry 2020, 32, 133–151.

(4) James, E. I.; Murphree, T. A.; Vorauer, C.; Engen, J. R.; Guttman, M. Advances in hydrogen/deuterium exchange mass spectrometry and the pursuit of challenging biological systems. Chemical reviews 2021, 122, 7562–7623.

(5) Hvidt, A.; Nielsen, S. O. Hydrogen exchange in proteins. Advances in protein chemistry 1966, 21, 287–386.

(6) Bai, Y.; Milne, J. S.; Mayne, L.; Englander, S. W. Primary structure effects on peptide group hydrogen exchange. Proteins: Structure, Function, and Bioinformatics 1993, 17, 75–86.

(7) Connelly, G. P.; Bai, Y.; Jeng, M.-F.; Englander, S. W. Isotope effects in peptide group hydrogen exchange. Proteins: Structure, Function, and Bioinformatics 1993, 17, 87–92.

(8) Nguyen, D.; Mayne, L.; Phillips, M. C.; Walter Englander, S. Reference parameters for protein hydrogen exchange rates. Journal of the American Society for Mass Spectrometry 2018, 29, 1936–1939.

(9) Hilser, V. J.; Freire, E. Structure-based calculation of the equilibrium folding pathway of proteins. Correlation with hydrogen exchange protection factors. Journal of molecular biology 1996, 262, 756–772.

(10) Vendruscolo, M.; Paci, E.; Dobson, C. M.; Karplus, M. Rare fluctuations of native proteins sampled by equilibrium hydrogen exchange. Journal of the American Chemical Society 2003, 125, 15686–15687.

(11) Best, R. B.; Vendruscolo, M. Structural interpretation of hydrogen exchange protection factors in proteins: characterization of the native state fluctuations of CI2. Structure 2006, 14, 97–106.

(12) Bradshaw, R. T.; Marinelli, F.; Faraldo-Gómez, J. D.; Forrest, L. R. Interpretation of HDX data by maximum-entropy reweighting of simulated structural ensembles. Biophysical Journal 2020, 118, 1649–1664.

(13) Nguyen, T. T.; Marzolf, D. R.; Seffernick, J. T.; Heinze, S.; Lindert, S. Protein structure prediction using residue-resolved protection factors from hydrogen-deuterium exchange NMR. Structure 2022, 30, 313–320.

(14) Devaurs, D.; Antunes, D. A.; Borysik, A. J. Computational modeling of molecular structures guided by hydrogen-exchange data. Journal of the American Society for Mass Spectrometry 2022, 33, 215–237.

(15) Stofella, M.; Grimaldi, A.; Smit, J. H.; Claesen, J.; Paci, E.; Sobott, F. Computational Tools for Hydrogen–Deuterium Exchange Mass Spectrometry Data Analysis. Chemical Reviews 2024, 124, 12242–12263.

(16) Gold, V. Advances in Physical Organic Chemistry ; Elsevier, 1969; Vol. 7; pp 259–331.

(17) Schowen, K. B.; Schowen, R. L. [29] Solvent isotope effects on enzyme systems. Methods in enzymology 1982, 87, 551–606.

(18) Loh, S. N.; Markley, J. L. Hydrogen bonding in proteins as studied by amide hydrogen D/H fractionation factors: application to staphylococcal nuclease. Biochemistry 1994, 33, 1029–1036.

(19) Bowers, P. M.; Klevit, R. E. Hydrogen bonding and equilibrium isotope enrichment in histidine-containing proteins. Nature Structural Biology 1996, 3, 522–531.

(20) LiWang, A. C.; Bax, A. Equilibrium protium/deuterium fractionation of backbone amides in U13C/15N labeled human ubiquitin by triple resonance NMR. Journal of the American Chemical Society 1996, 118, 12864–12865.

(21) Shi, Z.; Olson, C. A.; Kallenbach, N. R.; Sosnick, T. R. D/H amide isotope effect in model α-helical peptides. Journal of the American Chemical Society 2002, 124, 13994–13995.

(22) Hwang, T.-L.; Van Zijl, P. C.; Mori, S. Accurate quantitation of water-amide proton exchange rates using the phase-modulated CLEAN chemical EXchange (CLEANEXPM) approach with a Fast-HSQC (FHSQC) detection scheme. Journal of biomolecular NMR 1998, 11, 221–226.

(23) Chevelkov, V.; Xue, Y.; Krishna Rao, D.; Forman-Kay, J. D.; Skrynnikov, N. R. 15NH/D-SOLEXSY experiment for accurate measurement of amide solvent exchange rates: application to denatured drkN SH3. Journal of Biomolecular NMR 2010, 46, 227–244.

(24) Eigen, M. Proton transfer, acid-base catalysis, and enzymatic hydrolysis. Part I: elementary processes. Angewandte Chemie International Edition in English 1964, 3, 1–19.

(25) Zhou, G.-Q.; Zhong, W.-Z. Diffusion-Controlled Reactions of Enzymes: A Comparison between Chou’s Model and Alberty-Hammes-Eigen’s Model. European journal of biochemistry 1982, 128, 383–387.

(26) Molday, R.; Englander, S.; Kallen, R. Primary structure effects on peptide group hydrogen exchange. Biochemistry 1972, 11, 150–158.

(27) Mikkelsen, K.; Nielsen, S. O. Acidity measurements with the glass electrode in H2O-D2O mixtures. The Journal of Physical Chemistry 1960, 64, 632–637.

(28) Abel, E.; Bratu, E.; Redlich, O. Die elektrolytische Dissoziation des schweren Wassers. Zeitschrift für Physikalische Chemie 1935, 173, 353–364.

(29) Shoesmith, D. W.; Lee, W. The ionization constant of heavy water (D2O) in the temperature range 298 to 523 K. Canadian Journal of Chemistry 1976, 54, 3553–3558.

(30) Gold, V.; Lowe, B. Measurement of solvent isotope effects with the glass electrode. Part The ionic product of D2O and D2O–H2O mixtures. Journal of the Chemical Society A: Inorganic, Physical, Theoretical 1967, 936–943.

(31) Salomaa, P.; Schaleger, L. L.; Long, F. Solvent deuterium isotope effects on acid-base equilibria. Journal of the American Chemical Society 1964, 86, 1–7.

(32) Sweeton, F. H.; Mesmer, R.; Baes, C. Acidity measurements at elevated temperatures. VII. Dissociation of water. Journal of Solution Chemistry 1974, 3, 191–214.

(33) Gold, V.; Grist, S. Deuterium solvent isotope effects on reactions involving the aqueous hydroxide ion. Journal of the Chemical Society, Perkin Transactions 1972, 2, 89–95.

(34) Kiefer, P. M.; Hynes, J. T. Kinetic isotope effects for adiabatic proton transfer reactions in a polar environment. The Journal of Physical Chemistry A 2003, 107, 9022–9039.

(35) Stofella, M.; Seetaloo, N.; St John, A. N.; Paci, E.; Phillips, J. J.; Sobott, F. Recalibrating Protection Factors Using Millisecond Hydrogen/Deuterium Exchange Mass Spectrometry. Analytical Chemistry 2025, 97, 2648–2657.

(36) Grimaldi, A.; Stofella, M.; Hobbs, B.; Karamanos, T. K.; Paci, E. Amide Hydrogen Deuterium Exchange in Isotopically Mixed Waters. arXiv preprint 2510.24860 2025,

(37) Van Rossum, G.; Drake Jr, F. L. Python reference manual ; Centrum voor Wiskunde en Informatica Amsterdam, 1995.

